# Spatially distinct cellular and molecular landscapes define prognosis in triple negative breast cancer

**DOI:** 10.1101/2025.02.10.637503

**Authors:** Kavitha Mukund, Darya Veraksa, David Frankhouser, Lixin Yang, Jerneja Tomsic, Raju Pillai, Srijan Atti, Zahra Mesrizadeh, Daniel Schmolze, Xiao-Cheng Wu, Mary-Anne LeBlanc, Lucio Miele, Augusto Ochoa, Victoria Seewaldt, Shankar Subramaniam

## Abstract

**Background:** Triple-negative breast cancer is a prevalent breast cancer subtype with the lowest 5-year survival. Several factors contribute to its treatment response, but the inherent molecular and cellular tumor heterogeneity are increasingly acknowledged as crucial determinants.

**Methods:** Spatial transcriptomic profiling was performed on FFPE tissues from a retrospective, treatment-naive group of women with differential prognoses (17 with >15 years survival-good prognosis (GPx) and 15 with <3 years survival-poor prognosis (PPx)) using GeoMX^®^ Digital Spatial Profiler. Regions of interest were segmented on pan-cytokeratin and analyzed for tumor and stromal components, probed using GeoMx human whole transcriptome atlas (WTA) panel. Data quality control, normalization, and differential analysis was performed in R using GeomxTools and linear mixed models. Additional analyses including cell-type deconvolution, spatial entropy, functional enrichment, TF-target / ligand-receptor analysis and convolution neural networks were employed to identify significant gene signatures contributing to differential prognosis.

**Results:** Here we report on the spatial and molecular heterogeneity underlying differential prognosis. We observe that the state of the epithelia and its microenvironment (TME) are transcriptionally distinct between the two groups. Invasive epithelia in GPx show a significant increase in immune transcripts with the TME exhibiting increased immune cell presence (via IF), while in PPx they are more metabolically and translationally active, with the TME being more mesenchymal/fibrotic. Specifically, pre-cancerous epithelia in PPx display a prescience of aggressiveness as evidenced by increased EMT-signaling. We identify distinct epithelial gene signatures for PPx and GPx, that can, with high accuracy, classify samples at the time of diagnosis and likely inform therapy.

**Conclusions:** To the best of our knowledge, this is the first study to leverage spatial transcriptomics for an in-depth delineation of the cellular and molecular underpinnings of differential prognosis in TNBC. Our study highlights the potential of spatial transcriptomics to not only uncover the molecular drivers of differential prognosis in TNBC but also to pave the way for precision diagnostics and tailored therapeutic strategies, transforming the clinical landscape for this aggressive breast cancer subtype.

## INTRODUCTION

Triple-negative breast cancer (TNBC) is a highly aggressive breast cancer (BC) subtype, characterized by <1% expression of estrogen and progesterone receptors (ER and PR) and wild-type expression of HER2/neu. While TNBC generally carries a poor prognosis[1], not all TNBCs are fatal. Treatment efficacy, prognosis, and subsequent survival are increasingly acknowledged to be affected by interactions of the tumor cells with its surrounding cellular milieu (tumor microenvironment, TME)[2,3]. High-throughput studies over the past decade have revealed the molecular and cellular heterogeneity of TNBC with implications for therapy[4–7]. Primarily based on bulk gene expression data, these studies highlight inter-patient heterogeneity and tumor immunogenicity but offer limited insights into specific interactions within the tumor ecosystem including molecular interactions between the malignant cells, stromal components (e.g., fibroblasts, adipocytes, pericytes, immune cells), vasculature, or the influence of environmental factors like hypoxia, stress, or extent of immune infiltration. Recent advancements in single-cell and spatial technologies address the limitations of bulk profiling technologies and offer increased granularity of the complex multidimensional interactions in tissue[8–16]. In particular, high-throughput spatial transcriptomic (ST) technologies such as GeoMX^®^ Digital Spatial Profiler (DSP), spotted ST arrays, and Visium 10X allows researchers to uniquely characterize the molecular and cellular basis of these complex interactions while conserving the tissue architecture (context specificity), in a high-throughput manner[13–15,17]. Though the use of ST in deconvoluting TNBC is in its infancy, studies have begun to provide novel insights. For instance, Wang et al[13] in their preprint, leverage previously published Lehmann molecular classification to highlight intra-tumoral spatial heterogeneity in 92 TNBC samples using spotted ST arrays. They show the differential contribution of the tumor and TME to the molecular subtypes and identify the presence of 14 major combinations of the molecular subtypes (9 major “ecotypes”) in TNBC. Using Visium 10X, Bassiouni et al[14] resolve the transcriptomic state in a racially diverse TNBC cohort (28 samples, 14 women) and reveal consistent patterns of spatial exclusion and dependency in TNBC, with hypoxia and immune expression distinct across races. These studies provide a broad understanding of the cellular architecture of the TNBC tumor ecosystem while leveraging previously published molecular classifications to provide insights into the molecular underpinnings of the spatially resolved sections. In contrast to the approaches outlined above, we propose using spatial transcriptomics to identify regions of interest (ROIs) with epithelial cells in distinct pathological states and establish functional states for these epithelia and their surrounding TME. By applying it to a retrospective group of treatment-naïve women with differential prognoses, we seek to gain deeper insights into the tumor transcriptional landscape informing prognoses and survival. Further, we hypothesize that the aggressive biology that dictates subsequent survival and response to therapy in TNBC is, to some extent, hardwired into the genome and can be detected in the transcriptional state of the cellular landscape. This study, to our knowledge, represents the first attempt to systematically resolve the spatial landscape and molecular underpinnings of TNBC with differential prognoses.

In this study, samples from two retrospective treatment-naïve groups with 17 good prognosis women with >15 yr. overall survival (henceforth referred to as GPx samples), and 15 poor prognosis women with <3 yr. overall survival (PPx) were acquired at the time of diagnosis and analyzed using GeoMX^®^ DSP. We capture over 700 ROIs across different spatial regions of the tumor, with varying immune infiltration patterns and tumor epithelia in diverse pathological states along the continuum of tumorigenesis. We further segment the ROIs on cytokeratin 15 (CK15) to independently investigate the epithelia and surrounding cellular milieu. We develop and adopt a unique annotation schema to consistently categorize the ROIs (Extended Data Fig 1a) on the histomorphology and design our downstream analysis. Our results show that the epithelia are transcriptionally distinct between the GPx and PPx groups - GPx (invasive) epithelia show a significant increase in immune transcript expression, while PPx (invasive) epithelia are more metabolically and translationally active, indicative of rapid disease progression. PPx epithelia show a transcriptionally prominent transition from pre-invasive states to invasiveness with activation of pathways associated with tumor progression, alluding to the prescience of aggressiveness in the PPx pre-invasive epithelia. Our analyses further emphasize that the cellular composition of the TME is distinct in GPx and PPx women, with GPx exhibiting an increased immune cell presence and the PPx displaying a more mesenchymal-/fibrotic and presence of exhausted CD8 T cells in their TME. We finally learn (using deep learning) a distinct epithelial multi-gene signature that can, with high accuracy classify samples as GPx or PPx, at the time of diagnosis and inform therapy (Fig. 1a).

**Figure 1.**
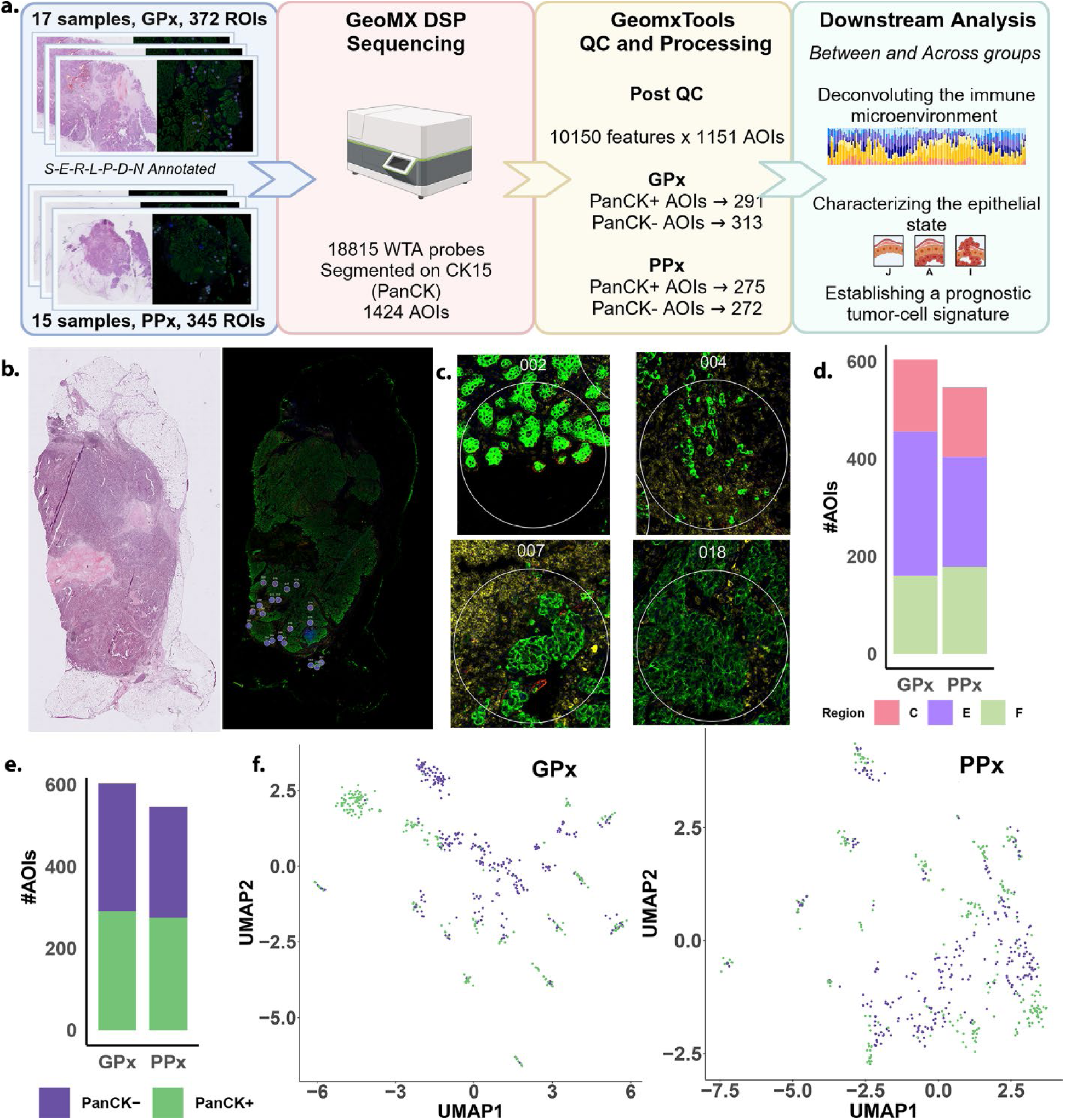
Study design and processing: **a.** Schematic of the study design and processing is presented. 17 good prognosis (GPx) and 15 poor prognosis (PPx) TNBC samples were selected, and H&E stained, with regions of interest (ROIs, purple dots show in IF images) carefully selected using a standardized approach. 4 channels/antibody stains were utilized to visualize the cells in IF (PanCK+ green, CD3+ yellow, CD45+ red, DAPI blue). Selected ROIs were subject to GeoMX profiling, and the .DCC files subsequently generated was QC-ed and processed using GeomxTools. Q3 normalized data was utilized for downstream processing and analysis, unless otherwise stated. b. Exemplar H&E slide with the IF imaging c. Exemplar ROIs on the slide in b are shown here. ROIs were selected by pathologist/trained clinician at the tumor interface, within the center of the tumor (with or without immune cells), and if available, regions distant from the center and edge (isolated foci). If available, regions with morphologically normal and/or precancerous changes (PanCK+), and with varying immune infiltration patterns (CD3+, Cd45+) regions were also chosen for ROIs. d. Barplot showing the distribution of post processed AOIs captured from different annotated regions of the slides) e. Barplot summarizing the distribution of the post processed AOIs, showing a fairly equal number of PanCK+ and PanCK- segments (604 GPx and 547 PPx AOIs)f. UMAP of AOIs from all the GPx and PPx samples post processing, showing fair separation between the segments, indicative of difference in underlying biology of the cellular landscape captured by segmenting, across a majority of the ROIs in both GPx and PPx.

## RESULTS

### Study design and development of a ROI annotation strategy

Utilizing GeoMX^®^ DSP we spatially profiled the cellular heterogeneity in 32 TNBC samples with documented differential prognoses (retrospective). TNBC samples were defined by <1% protein expression of ER and PR by immunohistochemistry (IHC) and no evidence of HER2/neu gene overexpression or amplification by combining IHC and FISH. All samples were reviewed by a single dedicated pathologist (D.S.) FFPE breast tissues of 17 treatment-naïve women GPx, and 15 treatment-naïve PPx women were chosen from the Louisiana Tumor Registry (IRB#1169). In GPx, 8 women self-identified race as Black, and 9 as White, while in PPx 5 women self-identified as Black and 10 as White. The samples were H&E stained as well as subject to immunofluorescence (IF) imaging using 4 IF antibodies to identify various cell types including DAPI (blue) to identify nuclei, CD3 (yellow) for T-cells, CD45 (red) pan-immune cells, and pan-cytokeratin, CK15 (green) for epithelial cells (e.g. in Fig. 1b). Regions of illumination/interest (ROIs, diameter=300 microns) were selected by pathologist/trained clinician at the tumor interface, within the center of the tumor (with or without immune cells), and when available, regions distant from the center and edge (isolated foci). Regions with morphologically normal and/or precancerous changes in the epithelial cells were also selected as ROIs. GeoMX additionally offers the ability to segment areas of illumination/interest (AOIs) within ROIs using fluorescently labeled antibodies to independently transcriptionally profile each segmented area. We segmented each ROI on the cytokeratin antibody (CK15) to identify epithelial cells within each ROI, henceforth referred to as the PanCK+ AOI/segment, and all the negative space captured with the ROI as PanCK- AOI/segment representing the surrounding tumor milieu. Exemplar ROIs with varying immune infiltration, and pathological epithelial states, taken from different regions of the excisions are shown in Fig. 1c.

We identified a total of 717 regions of interest (ROIs, 345 PPx and 372 GPx) and captured a total of 1424 AOIs (737 GPx and 687 PPx, Fig. 1d) from the 32 samples, with uniform distribution of segments across samples (Fig. S1a). To meaningfully, consistently and systematically document and categorize the ROIs within the slides and across the samples, we developed and adopted a unique annotation schema and used this in our downstream analysis. Each of the 717 ROIs were annotated using the S-E-R-P-L-D-N annotation schema. Briefly, each ROI was annotated by a pathologist/clinician from IF images, with the following descriptors: Sample (S), Region (R), Epithelial type (E), Immune cell localization (L), Immune cell population (P), Immune cell distribution (D) and Immune cell number (N) as detailed in Supplementary Table 1. These descriptors allowed us to uniquely classify each ROI for downstream algorithmic processing. We probed the 717 ROIs/1424 AOIs using GeoMX Whole Transcriptomic Atlas (GeoMX Hu WTA™), containing 18815 probes. The 1424 DCC files generated were subsequently processed using the GeomxTools[18] (see Methods for details). Probe and gene QC (see Methods) of the 1424 AOIs resulted in a filtered data object containing 10150 genes across 1151 AOIs (604 GPx and 547 PPx, Fig. 1e, Extended Data Fig 1b). Extended Data Figs. 1c and d summarize the distribution of ROIs pre- and post-QC, from different regions of the slide (center, edge and isolated foci), as well as AOIs with differing epithelial morphology (Invasive I, and others such as A, D, J, N, and O). Specifically, given their distribution, we only emphasize ROIs that contain epithelial cells that exhibit atypia (A); invasiveness/transformed (I) and appear normal (adjoining areas of atypia/invasiveness (J)) to delineate the state of the tumor ecosystem and interactions with the surrounding TME.

Most of the ROIs from GPx and PPx samples, post-processing, showed separation between the segments (AOIs), indicative of differences in the underlying biology of the cellular landscape captured by segmenting, (Fig. 1f). Clustering of the pooled signal from each ROI (average expression across PanCK+ and PanCK- AOIs) captured the inter-patient and intra-tumoral heterogeneity of the TNBC set (Fig. S1e). To better dissect this heterogeneity, we characterized the molecular underpinnings and state of the tumor and its TME as captured by the PanCK- and PanCK+ segments and the interactions that exist across them described in the following sections.

### Deconvoluting the cell-type repertoire of the tumor microenvironment in good and poor prognosis TNBC

Spatial segmentation of each ROI on PanCK captured the non-epithelial cell milieu in the PanCK- region, including stromal cells and the tumor-infiltrating lymphocytes (TILs), representative of the TME in our TNBC samples. Specifically, the 313 GPx and 272 PPx PanCK- AOIs captured showed varying levels of immune infiltration as detected using the CD3+ (yellow channel) and CD45+ (red channel) via immunofluorescence imaging. Intensity-based immune quantification (red channel) showed a significant difference between GPx and PPx(p<0.05, albeit the yellow channel being statistically non-significant) (Fig. 2a, b). Recent research has suggested a differential immune cell distribution between tumor edge and center may influence tumor aggressiveness[34]and prognosis in other cancers. We observed prominent transcriptional dysregulation in the TME between AOIs from center to edge in both GPx (albeit not statistically significant, Fig. S2a) and PPx (Fig. 2c, Supplementary Table 2). In PPx, however, the TME of AOIs from the center (vs edge) showed a distinct increase (adj p <0.05) in genes associated with metabolic and cellular stress and remodeling including ENO1, CYC1, ATP5F1C, HSPA1B, and AHSA1. Increased expression of genes associated with tumor survival and immune evasion was seen at the edge (vs center) including genes such as NR3C1, BCL2, and AP3BP1. We observed no statistically (fdr<0.05) significant difference between the TME of GPx and PPx, chosen at different regions of the tumor slide-Center, edge and isolated foci (Fig. S2b). A comparison of the genes expressed in the PanCK- segments irrespective of the spatial region between GPx and PPx (M-A plot, Fig. 2c) however showed a higher relative expression in GPx-TME for genes involved in immune response such as B2M, CD74, immunoglobulins and HLA-DRB1; while PPx-TME showed increased expression of genes associated with abnormal cell morphology and EMT including PTBP1, COL1A1, COL1A2, COL6A2, FLNA, MYH9, and VIM.

**Figure 2.**
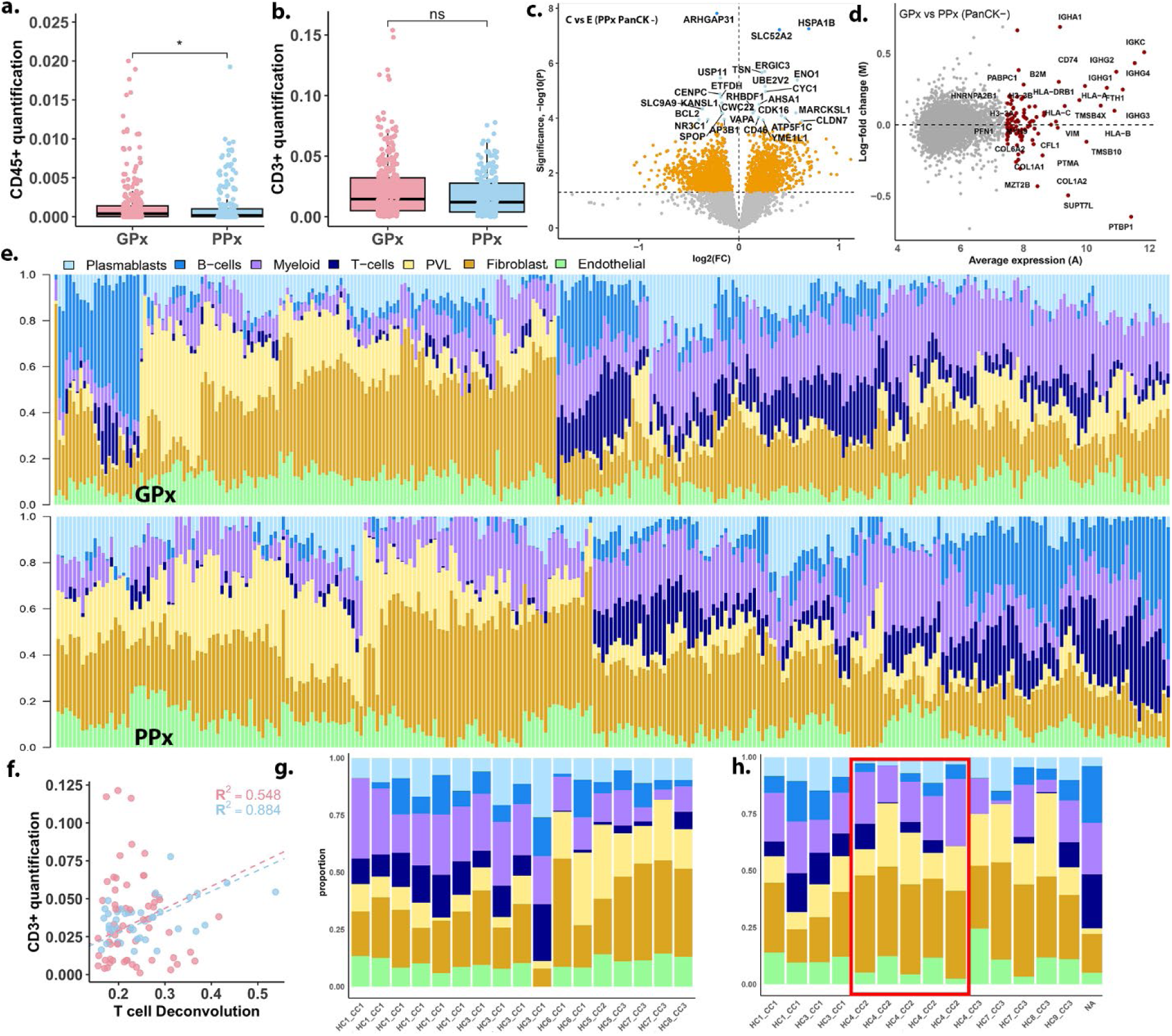
Immune deconvolution of the tumor microenvironment in good (GPx) and poor prognosis (PPx). a. and b. Quantification of CD45 and CD3 channel intensities in the PanCK- segments. c. Differentially expressed genes captured in the comparison of AOIs from center (C) to edge (E) from the PanCK- segments in PPx. Gene names are highlighted when p.adj < 0.05. d. log –fold change (M) to average expression (A) of GPx vs PPx PanCK- AOIs. The top 100 genes are highlighted in red, with the top 30 named on the plot. PPx shows a more distinct increase in expression of genes that are actively involved in EMT including PTMA, VIM, PTBP1 etc. e. The normalized frequency of immune cell type deconvoluted in 313 GPx (top panel) and 272 PPx (bottom panel) PanCK- AOIs generated using SpatialDecon and a custom reference profile matrix. We estimated the abundance of 7 major immune cell types (T-cells, B- cells, plasmablasts, myeloid, PVLs and fibroblast (mesenchymal cell types) and endothelial cells). f. Correlation between the CD3+ channel intensity captured and T cell deconvolution in the PanCK- segments, captured in the GPx (red) and PPx (blue) samples. g. Sample level estimation of deconvoluted cell frequencies for GPx, with the names corresponding the subtype identified in our earlier study. h. Similarly, for PPx. The red box highlights the reduced presence on T-cells by deconvolution in women classified as CC2.

To gain quantitative insights into the differential contribution of various cell types to the TME in GPx and PPx TNBC, we utilized SpatialDecon[35] v1.1. As outlined in Methods, we first generated a custom reference profile matrix of TNBC (see Methods for details), for use in SpatialDecon. Utilizing this, we estimated the abundances of 7 major cell types (T-cells, B- cells, plasmablasts, myeloid, mesenchymal cells (perivascular-like (PVLs) and fibroblasts) and endothelial cells) for both GPx and PPx samples (Fig. 2d). Correlation of the deconvoluted T-cell proportions with the proportion of T-cells captured via IF (CD3+ channel intensities in GPx and PPx (average r^2^∼0.7, Fig. 2e, see Methods) showed good concordance. A visual inspection of the abundances (Fig. 2d) highlighted: 1. the presence of a richer immune repertoire (T, B, and plasmablasts) within GPx, and a richer myeloid cell presence, in AOIs that showed immune infiltration; 2. higher mesenchymal content (fibroblast and PVL) within PPx. Based on the analysis of quantile distributions (IQR and range) of cell type frequencies we found that GPx AOIs exhibit a slightly more varied and richer immune presence (myeloid cells, T, B, and plasmablasts), consistent with the visual summary (Fig. S2c, Supplementary Table 3). A similar analysis of PPx AOIs likewise showed a stronger and more varied mesenchymal presence (PVL+Fibroblasts) compared to GPx. Our group had previously subtyped 31 of the 32 patients profiled here[7] into three distinct groups (CC1- increased immune infiltration; CC2- immune desert and CC3- increased fatty acid/nuclear receptor signaling) using bulk-expression measurements Interestingly, a higher proportion of patients in the CC2 and CC3 subgroups were identified within our PPx (∼66% of the samples). Furthermore, utilizing the spatial deconvolution calculated above, we averaged frequencies across all AOIs for each patient in GPx (Fig. 2g) and PPx (Fig. 2f) (see Methods for details). We observed that PPx women subtyped into the CC2 subgroup showed a distinct lack of T-cells in the TME, consistent with the definition of CC2 (red box, Fig. 2f). GPx had a higher proportion of samples belonging to the CC1 subtype (∼64%), which we have previously shown to correlate with better prognosis. Given the differential T-cell distribution between the prognostic groups, we examined the nature of the T-cell enriched AOIs in the following section.

### GPx and PPx TNBC exhibit distinctly different cell states in the TME with significant immune infiltration

Recent research suggests that only a small portion of the TILs exist in a tumor-reactive state, implying that a majority of TILs are quiescent and not functional[36]. We asked if there was a difference in the functional state of the TILs and surrounding stroma, within ROIs with significant T-cell infiltration identified by deconvolution, across GPx and PPx, irrespective of the epithelial state captured within the AOI (A, J, or I). To control for the intra-patient heterogeneity, we chose patient samples that showed greater than a third of their AOIs with a significant proportion of T-cells (>0.75 quantiles) confirmed via IF, resulting in 61 AOIs from 5GPx women (Fig. S2c) and 40 AOIs from 3 PPx women (Fig. S2d) and, henceforth referred to as the T_enr_ subsets. To investigate the spatial organization of the immune cells in these T_enr_ subsets across GPx and PPx, we performed a spatial entropy analysis on immunofluorescent images of these regions. We calculated the spatial partial information (SPI), a measure of how much spatial organization can explain the distribution of staining. In the GPx-T_enr_ subset, CD3+ staining showed significantly higher SPI with PanCK+ staining at distances of ∼28 and 70μm (Fig. 3a). The distance here signifies the average pairwise distance from the distribution profile. A higher SPI implies that the distribution of T-cells is more dependent on tumor cell organization. We see a higher SPI in GPx- T_enr_ than PPx-T_enr_ implying increased interaction of T-cells with tumor cells in the GPx-T_enr_ subset.

**Figure 3.**
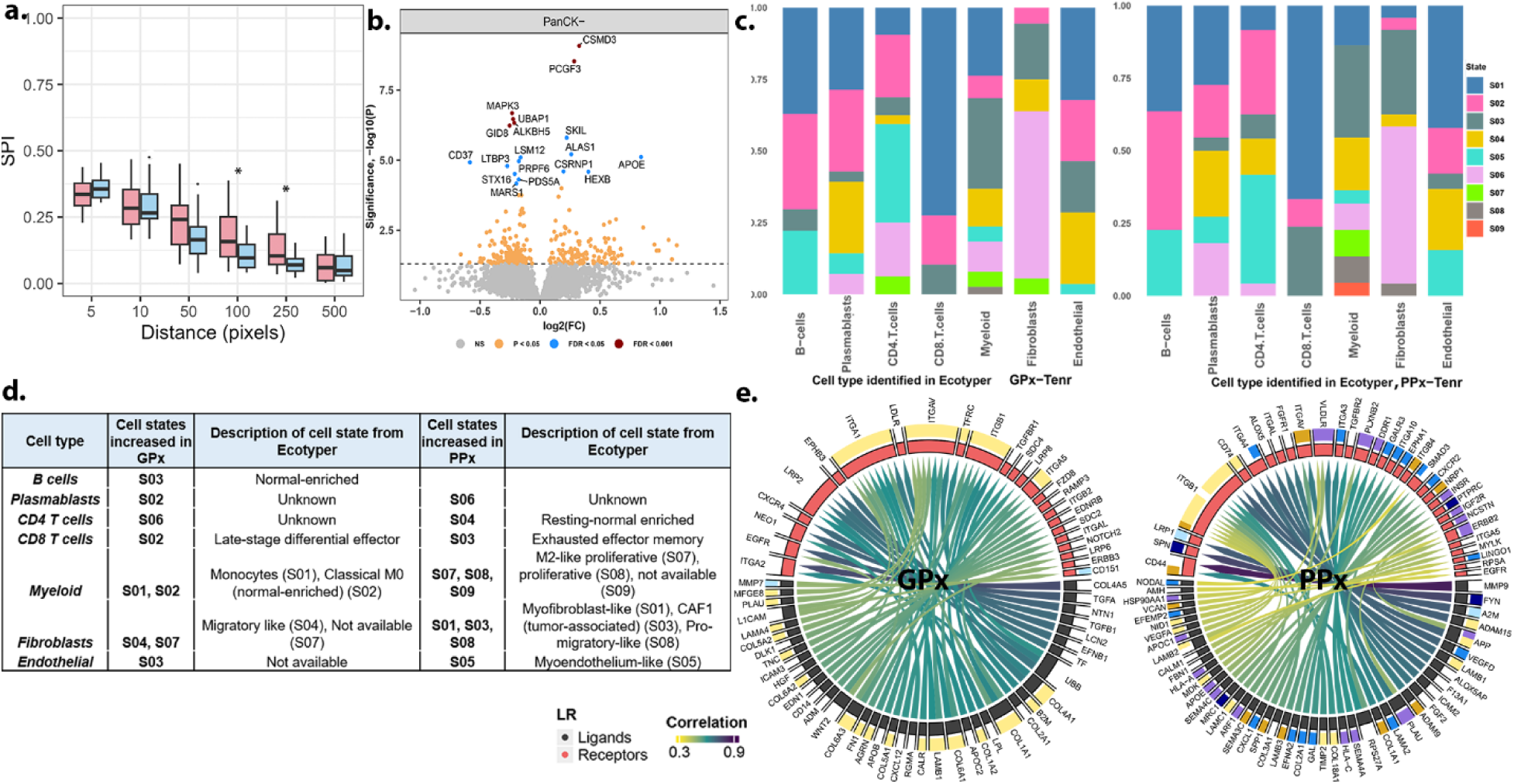
Establishing the transcriptional states of immune cells in the T-enr subset. a. SPI spatial partial information (SPI) (measure of spatial entropy) calculated for CD3+ vs PanCK+ channels in GPx-Tenr (red) and PPx-Tenr (blue). b. The differentially expressed genes identified by comparing the expression of GPx and PPx PanCK- AOIs. c. Normalized distribution of cell states identified using EcoTyper in AOIs that were enriched for immune (T) cell infiltration within GPx and h. within PPx. d. A table summarizing the cell states that are preferentially increased within GPx-Tenr and PPx-Tenr as captured in c. e. A chord diagram highlighting the top 50 unique ligand-receptor (L-R) interactions (at corr>0.25, q<0.05) identified using BulkSignalR, within the TME of the GPx- Tenr (left) and PPx- Tenr (right), respectively.

Differential analysis of genes expressed in the PanCK- segments of GPx-T_enr_ w.r.t PPx-T_enr_ (at a restrictive threshold of adj p <0.05, see Methods), showed significant upregulation of genes such as CD37, MAPK3, MARS1 and GID8 in PPx, that have been previously shown to correlate with poor prognosis (Fig. 3b). However, these measurements (and results) provide a unique challenge to discretizing the “transcriptomic state” of the various deconvoluted immune cell types, as the measurements are made on tens to hundreds of nuclei and across cell types within the PanCK- segments. To overcome this challenge, we adopted a manifold approach to characterizing the transcriptional states for the deconvoluted cell types in GPx and PPx. We first leveraged and adapted a recently published machine learning framework called EcoTyper[26], which defines distinct *functional cell states* for various cell types in carcinomas, using publicly available bulk expression data (see Methods). Using EcoTyper, and the gene expression measured within the PanCK- segments, we identified that a majority of the cell types exist in similar cell states within both GPx and PPx (Fig. 3c, See Methods). For instance, most of the fibroblasts exist in state S05, while B cells exist in naïve (S01) and activated (S02 and S05) states. Notably, however, there are certain cell states more pronounced within PPx-T_enr_ (tabulated in Fig. 3d). For instance, fibroblasts in PPx-T_enr_ showed increased presence of myofibroblast-like (S01), CAF1 (S03), and pro-migratory states (S08). A higher proportion of AOIs were found to be in the CD8 T-exhausted state (S03) within PPx- T_enr_. Additionally, we observed a significant difference in the average expression of cell-state markers (see Methods) for several of these unique states for instance, CD8 T cells in S03 in PPx had increased expression compared to AOIs in S03 within GPx (Fig. S2e). Next, utilizing BulkSignalR[27] we identified 123 and 69 ligand-receptor (L-R) interactions (at corr>0.25, q<0.05, see Methods) that are uniquely exhibited in the TME of the GPx- T_enr_ and PPx- T_enr_, respectively (Fig. 3e, Supplementary Table 4). Cell types that were most correlated with each LR-Interactions based on the deconvoluted abundances were also computed. Specifically, several of the L-R interactions uniquely identified within the PPx-T_enr_ stroma such as COL1A1-CD44, MMP9-CD44, and SPP1-ITGAV (Fig. 3e) are known to correlate with poor prognosis, a more mesenchymal tumor characterized by aggressive biology[37–39] of tumors, consistent with EcoTyper results.

### Characterizing the epithelial cell states in good prognosis and poor prognosis TNBC

The PanCK+ segments captured via GeoMX DSP allowed us unprecedented insights into the state of the epithelial cells within GPx and PPx. Broad characterization of the cell states was first performed using single-sample GSEA[40] (ssGSEA, see Methods) on 291 GPx and 275 PPx epithelial AOIs (PanCK+) (Fig. S3). ssGSEA results highlighted a prominent increase in mechanisms associated with proliferation, DNA repair and metabolism within the invasive epithelia, in both GPx and PPx (Fig. S3). The AOIs with non-invasive epithelia in both subsets were more prominently associated with signaling cascades involved in EMT including WNT β- catenin, Notch, and KRAS signaling. We also observed that a majority of the AOIs with tumor cells (or invasive epithelia (I)) clustered distinctly from AOIs with other epithelial types, in both GPx and PPx, irrespective of the region.

To improve our signal-to-noise ratio and emphasize molecular mechanisms that are significantly differential between GPx and PPx epithelia, we utilized non-negative matrix factorization[41] (see Methods) to reduce the number of features. 1712 *consensus features* (across 291 GPx PanCK+ AOIs and 1677 *consensus features* across 275 PPx PanCK+ AOIs see Methods, Supplementary Table 5) were identified and used for all further downstream analyses of the epithelial state. In the following sections, we sought to answer three broad biological questions about the epithelial cell state. 1. Does the tumor cell state dictate prognosis and can we define a transcriptional program underlying annotated epithelial states? 2. Is there a prescience of aggressive biology in pre-invasive states? 3. Can we identify a distinct tumor epithelial cell signature underlying each prognostic group?

### A distinct transcriptional profile underlies invasive tumor epithelia with differential prognosis

Epithelial tumors are characterized by rapid growth and differentiation and exist in varying states across the tumor. We hypothesize that the trajectory followed by epithelial cells from a non-invasive state to an invasive state differs in groups with differential prognosis, and subsequently proscribe prognosis.

As a first step, we assessed if the state of the tumor cell (invasive epithelia) is informed by the spatial location of the chosen ROIs (center or edge) of the tumor (Fig. 4a, Fig. S4a). A comparison of tumor cells (AOIs) from the center of the tumor to those at the edge of the tumor showed no statistically significant differences (at fdr<0.05) in either GPx or PPx patients (Fig. 4b). These indicated that the transcriptional state of the tumor epithelia (I) is independent of the spatial context. Next, we identified transcriptional programs underlying the invasive (tumor) epithelial state by comparing them with normal adjacent epithelia in both GPx and PPx (Fig. 4c, see Methods). A significant upregulation of genes (FDR <0.05) was observed in invasive epithelia, irrespective of prognosis, and included several histones such as H3C15, H4C15, H1-5, H2AC19, H2AZ1, H3C7; genes associated with cytoskeletal stability and cell cycle control including TMSB10, STMN1, CDKN2A were also highly differentially regulated (Fig. 4c, Supplementary Table 6). A closer inspection of the upregulated DEGs in the invasive epithelia of PPx and GPx (PPx-I and GPx-I), showed a large degree of overlap (191, Fig. 4d), with the overlapping genes being enriched for proliferation (G2M checkpoint, MYC targets, E2F targets, mTORC1 signaling, Fig. S4b). This finding is suggestive of a highly dynamic cell state of invasive epithelia compared to its normal-adjacent counterparts.

**Figure 4.**
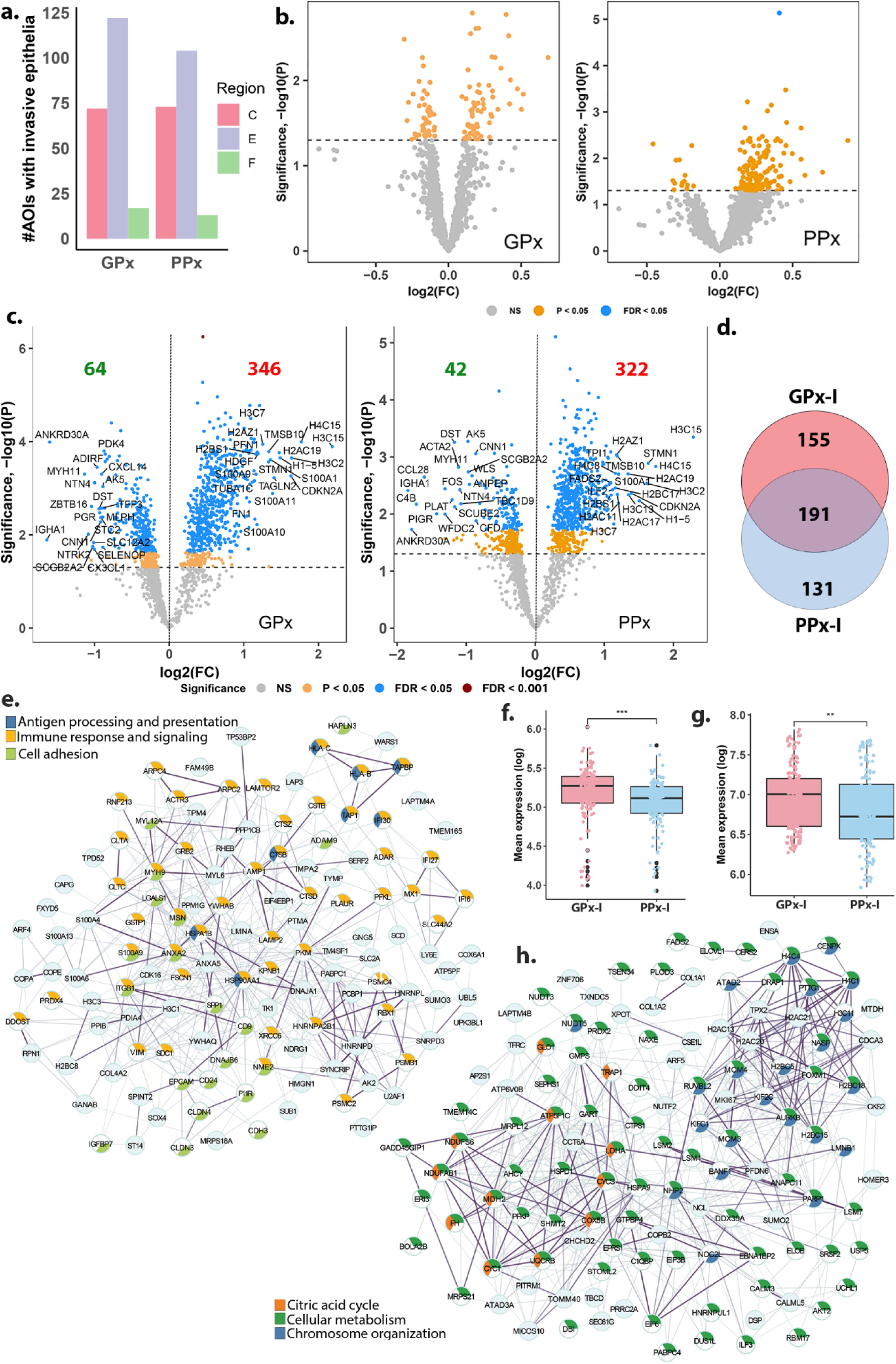
Characterizing the tumor cell state. a. Distribution of AOIs with invasive epithelia that were captured in the center (C), edge (E) and isolated foci (F) of the tumor samples in GPx and PPx. b. A comparison of consensus gene features in tumor cells (AOIs with invasive epithelia) from center of the tumor to those at the edge of the tumor showed no statistically significant differences (at FDR<0.05) in either GPx or PPx patients. c. Differential gene expression identified significant transcriptional dysregulation in invasive epithelia compared to normal-adjacent epithelia from GPx (204 AOIs, left) and PPx (184 AOIs, right). d. A Venn highlighting the overlap in genes upregulated within the invasive epithelia from GPx (GPx-I) and PPx (PPx-I). e. Protein-protein interaction network generated using the uniquely dysregulated genes GPx-I (155/346) form a tightly connected network of interacting genes PPI enrichment adj p <10-16) associated with immune signaling, antigen processing and presentation (APC), and cellular adhesion including genes such as MX1, IFI6, CTSB/D, HSP90AA1, PSMB1, PSMC4, PSMC2, ANXA2/5. f. Box plot of the mean expression of 54 immune genes identified within the PPI shows a significant increase in GPx- I AOIs compared to the PPx-I AOIs. g. Mean expression of the recently published optimized immune response signature (oIRS) in GPx-I and PPx-I, shows a statistically significant preferential increase in the tumor epithelia of GPx-I compared to PPx-I h. A PPI of uniquely upregulated genes in PPx-I showed evidence for extensive metabolic reprogramming and chromatin remodeling including genes such as MKI67, NDUFAB1, NDUFS6, HSPA9, AKT2, PFKP, PRDG2, MDH1/2, ATP5F1C, CTPS1, LDHA (mitochondrial and cellular metabolism, fdr<0.001) and AURKB, KIF2C, KIF1C, H2As, TPX2 (chromatin remodeling/organization, fdr<0.001

Interestingly, however, DEGs uniquely dysregulated within each prognostic group, compared to their normal adjacent showed activation of distinct programs. Notably, uniquely differentially expressed genes in the GPx invasive epithelia (155/346) were identified to belong to a tightly connected network of interacting genes (PPI enrichment adj p <10^-16^, Fig. 4e, Fig. S4c) associated with immune signaling, antigen processing and presentation (APC), and cellular adhesion including genes such as MX1, IFI6, CTSB/D, HSP90AA1, PSMB1, PSMC4, PSMC2, ANXA2/5. Transcription factors (TFs) such as API1, FOS, and STAT3 were identified as TFs (using DecoupleR[31]) driving this response of the upregulated DEGs identified uniquely in GPx-I (Fig. S4d). STAT3 has been previously reported to correlate with favorable prognosis[42]. Furthermore, the 54 immune transcripts identified within this module showed significantly higher expression in GPx-I compared to PPx-I epithelia (Fig. 4f, Supplementary Table 7). These results are exciting in light of the recent research that has indicated immune mimicry of tumor epithelial cells is widespread in tumors and is portrayed as a feature distinct to oncogenic transformation[42,43]. Utilizing the recently published optimized immune response signature (oIRS, Supplementary Table 7) of tumor cells[42], we further observed a preferential increase of oIRS in the tumor epithelia of GPx-I compared to PPx-I (Fig. 4g). Several components also involved in APC were uniquely increased in GPx-I. APC has been previously suggested to be an independent prognostic factor in BC[42]. A similar analysis of the uniquely upregulated genes in PPx-I showed evidence for extensive metabolic reprogramming and chromatin remodeling including genes such as MKI67, NDUFAB1, NDUFS6, HSPA9, AKT2, PFKP, PRDG2, MDH1/2, ATP5F1C, CTPS1, LDHA (mitochondrial and cellular metabolism, fdr<0.001) and AURKB, KIF2C, KIF1C, H2As, TPX2 (chromatin remodeling/organization, fdr<0.001) (Fig. 4h, Fig. S4e). Transcription Factor (TF)-target enrichment analysis of the uniquely upregulated DEGs showed increased activity of TFs across a majority of the AOIs, such as HIF1A, MYC and AR that are known to be crucial determinants of oxidative, glycolytic and lipid metabolism[44–46] (Fig. S4f).

### Pre-invasive epithelia show evidence of aggressive biology in poor prognosis TNBC

Based on our earlier observation that the tumor epithelial (invasive) cell state is different between groups that display differential prognosis, we hypothesized that the aggressive biology associated with poor prognosis TNBC is more apparent in the pre-invasive epithelial state of the PPx samples. To test this hypothesis, we compared the transcriptional landscape of the pre-invasive state (A) to invasive epithelia within both groups (PPx and GPx). We observed that PPx exhibits a more dominant transcriptional regulation in transitioning from atypia to invasive epithelia (567 DEGs, fdr<0.05, Fig. 5a right), compared to GPx. Consistent with our hypothesis, we observed that the 134 genes upregulated in PPx-atypia (PPx-A) were largely associated with EMT (loss of baso-apical polarity), regulation of cell migration, cell motility and cell-cell adhesion (Supplementary Table 8, Fig. 5b). Increased expression of chemokines was observed in the atypia of PPx, including CXCL12, CXCL14 and CXCR2. PPx-A also shows an increased presence of KRT15 and KRT34 transcripts that have been suggested to correlate with an invasive phenotype in certain epithelial tumors with poor prognosis[47]. Evidence in the literature points to complement activation in tumor initiation and the role of tumor cells in modulating complement activation within the tumor microenvironment (TME)[48]. We noted that the expression of several upstream regulators in the classical complement cascade including C1QA/B/C, C1R, C1S, C2, C3, and C4B was significantly reduced in the transition from PPx-A to PPx-I (Fig. 5c). In contrast, they were notably higher in PPx-A compared to GPx-A (Fig. 5d, Fig. S5. To better characterize the role of complement in the interactions between tumor cells and the TME in GPx-A and PPx-A, we performed L-R analysis on PanCK+ and PanCK- in ROIs with atypic epithelia using BulkSignalR (see Methods). We identified 115 L-R interactions unique to GPx-A and 140 interactions unique to PPx-A (Fig. 5e, Supplementary Table 9). We noted that both groups showed distinct C3-associated interactions (correlated in epithelia)- PPx-A showed C3-CD81 and C3-LRP1 binding. Expression of CD81 (on B-cells and exosomes secreted by CAFs) and LRP1 (expressed on epithelial cells and fibroblasts) have both been previously associated with EMT and poor prognosis[49–51]. C3-CD81 and C3- LRP1 interactions captured here could indicate anaphylatoxin-mediated priming of precancerous epithelia into tumor cells capable of increased migration. In contrast, GPx-A was enriched for C3- ITGAX, likely indicative of an anti-tumorigenic opsonization event^39^.

**Figure 5.**
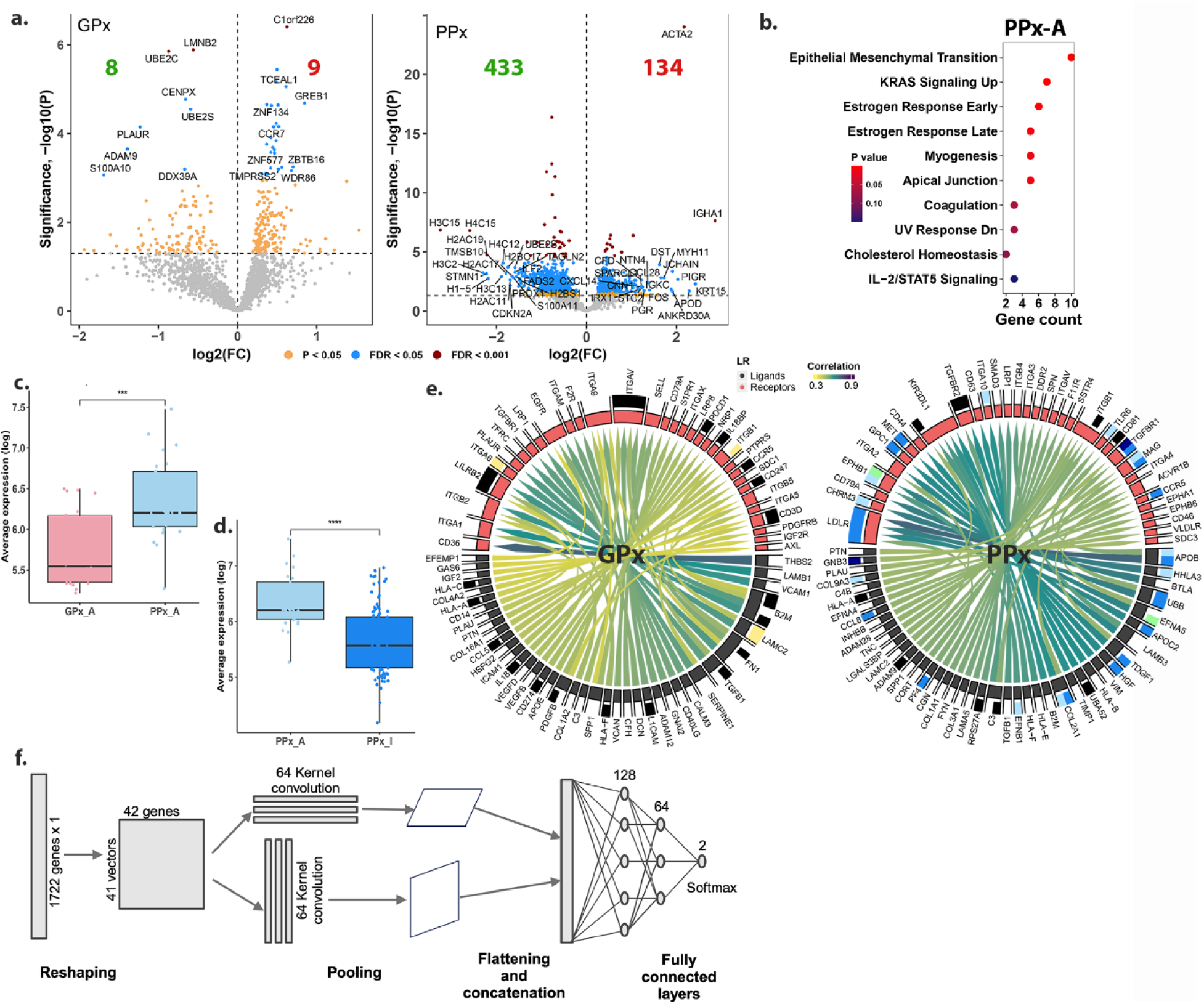
Characterizing the pre-invasive epithelial states and establishing a prognostic tumor cell signature. a. Differential expression analysis comparing the atypic to the invasive epithelia. PPx samples show a more dominant dysregulation of genes along this transition. b. Hallmark gene set enrichment of the genes identified as being upregulated in the atypia compared to invasive epithelia of PPX. c. Boxplots showing significantly higher expression of genes (mean expression) involved in the complement cascade within PPx-A compared to GPx-A AOIs. d. Boxplots showing significant increase in expression of genes in the complement cascade within AOIs with pre-invasive epithelial compared to AOIs with invasive epithelia (PPx-I) e. The L-R diagram highlighting the top 50 unique interactions captured within GPx and PPx atypia (with enrichment for epithelial cell types (black)). f. A schematic representation of the CNN model used for identification of epithelial gene signatures discriminatory of GPx and PPx to (235 and 236 genes respectively) that are outlined in Supplementary table XX.

### Identification of an epithelial gene signature underlying differential prognosis

Results presented in the sections above outline a distinct transition along the spectrum of epithelial states (J, A, I) in tumorigenesis. We further sought to identify a universal epithelial signature that is agnostic to cell state but dictates prognosis. To this extent, we re-ran NMF on the entire set of PanCK+ AOIs (566 AOIs, 291 GPx, and 275 PPx) and identified 1722 consensus features. We employed a deep-learning approach inspired by a previously published study[53], to identify an epithelial signature containing 235 and 236 genes that are essential to accurately label GPx and PPx epithelia (Supplementary Table 10). Briefly, the CNN developed within this study consisted of two distinct towers, with each tower, incorporating a Convolutional Neural Network (CNN) layer, followed by a LeakReLU activation function and a Max Pool layer with a kernel size of (2, 2) (Fig. 5f). The two pooled outputs were then flattened and concatenated before being fed into fully connected layers, initially with 128 nodes and ultimately narrowing down to 2 nodes, representing the model’s output “prognosis”. The model was trained using an Adagrad optimizer with a learning rate set at 0.001 and a Binary Cross-Entropy (BCE) loss function for a duration of 200 epochs. We generated discrete saliency maps for both good prognosis and poor prognosis to discern the most influential features. We next extracted genes that had a weightage of greater than 0.5, to ensure the importance of features in determining prognosis. The genes were then consolidated into a ranked list using Borda counts (see Methods). This process was repeated 100 times to ensure robustness and reliability. Results were only used from each newly trained model if its accuracy and precision on the test case were greater than an average of 90%. Finally, we compiled these ranked lists from each iteration into a comprehensive aggregated list, again using the Borda counts to establish a definitive ranking of features over these multiple iterations and represent the features that were most often used by CNN to predict the labels. A rank comparison of the two gene lists (see Methods) highlighted features/genes that were discriminatory (more dissimilarly ranked between the two lists) and were associated largely with a MYC-centric cellular stress response (red) and cell cycle control (purple) (Fig. S6) and included genes such as several histones H2BC11/17, H2AC13/19, MYC, YWHAZ, and CALR that have been previously shown to correlate with survival.

## DISCUSSION

The reciprocal interaction between the tumor and the microenvironment forms the basis for the observed intra-tumoral heterogeneity, impacting metastasis, prognoses and survival, and response to therapy in cancers[54]. Spatial transcriptomic profiling technologies have begun to offer unprecedented insights into characterizing this reciprocal interaction. To map the digital landscape of TNBC across tumor tissue from good and poor prognosis patient samples, we use the GeoMX spatial profiler. Further, given that the tumor landscape contains epithelial and stromal cells (constituting the TME) in a spatially heterogeneous distribution, here, we developed a novel ontological strategy to spatially annotate the cancer and TME in a biologically meaningful manner. This offers the unique feature of probing invasive and tumor-adjacent atypical epithelia toward understanding the causal mechanistic origins of tumor progression and prognosis. As we reported above, this spatial deconvolution enables us to traverse the tumor landscape along the spectrum of tumorigenesis delineating specific interactions of epithelial and surrounding stromal cells.

We sought to address whether the heterogeneity in the TME could aid in TNBC risk stratification. Despite the variability in the number of nuclei captured within each AOI in our digital landscape across samples, our analysis shows the increased presence of mesenchymal cell types (fibroblasts and PVLs) in PPx. Cell state deconvolution identified an increased presence of fibroblast states associated with poor survival (pro-migratory, myofibroblast, and CAF1). The presence of myofibroblasts is known to correlate with loss of epithelial differentiation, poor overall survival, and myeloid cell recruitment in various cancer types [55–57]. TNBC is widely recognized to exhibit increased immunogenicity compared to other BC subtypes[58]. CD8 T cells in several PPx- T_enr_ AOIs were captured to exist in an exhausted state, alluding to the role of immune exhaustion contributing to poorer prognosis[59]. A fibrotic TME was recently suggested to impede antitumor immunity by minimizing CD8^+^ T infiltration as well as mechano-metabolic reprogramming of TAMs in the TME[60] impacting subsequent prognosis. Furthermore, relative to measurements made in GPx, the presence of the states and ligand-receptor pairs and functional cell states routinely correlated with poor prognosis were evident in PanCK- segments of PPx samples. Although we observed no statistically significant difference in the transcriptional state of the TME between GPx and PPx TNBC, from different parts of the tumor tissue, the TME in PPx exhibited a distinct transcriptional profile depending on the locations within the tumor—center or edge— highlighting spatially contextual differences in metabolic activity and immune evasion strategies.

Phenotypic plasticity of tumor is largely dictated by the reciprocity of interactions and the environmental cues, between the TME and tumor epithelia and serve as a crucial determinant in tumor progression. This plasticity endows tumor epithelia with an ability to tans-differentiate, and de-differentiate (to a precursor or stem-cell-like state) and exhibits an EMT phenotype. Recent research in cancer highlights that this plasticity extends to the ability of epithelial cells to “mimic” other cell types[43,61]. In particular, “immune mimicry” of epithelial cells is suggested to facilitate the formation of a tumor immunosuppressive environment in pancreatic cancer and is an important feature of oncogenic transformation[62]. Furthermore, Donati et al[17] recently showed that in post-neoadjuvant chemotherapy TNBC patients with complete response to therapy spatial transcriptomics revealed an increased presence of immune activation markers in the PanCK+ regions of tumors. In this study, we generated a meta-gene signature to capture a reduced feature space from the epithelia of GPx and PPx. Utilizing this reduced feature space, we identified immune mimicry as a hallmark of invasive epithelia in treatment-naive GPx samples. Concomitantly, an increased immune presence was identified by both by transcript and spatial image quantification, within the TME of GPx samples. These findings are in concordance with recent research that has indicated an increased presence of TILs to correlate with better prognosis and therapeutic response[58,63]. Given our findings and extant research, we hypothesize that a preferential increase of immune transcripts in the invasive tumor epithelia is what primes the tumor towards a more immune-conducive TME and a better response to immunotherapy in GPx. We believe that this idea extends the notion of an “impressionable epithelium” and “dynamic reciprocity” indicative of the plasticity of epithelial cells in response to their surrounding milieu[64]

We examined the role of TME in dictating aggressive biology previously described as the “field effect hypothesis” [65]. The pre-invasive epithelia in PPx showed evidence of aggressive biology that included the increased expression of EMT-associated gene cascades and the presence of chemokines including CXCL14 (compared to invasive); expression of CXCL14 has been previously suggested to facilitate paracrine signaling between the tumor epithelia and TME to facilitate tumor aggressiveness in DCIS[66]. Differential early complement activation and signaling in PPx- compared to GPx-atypical epithelial cells (A). This may suggest an earlier and more dominant mesenchymal transition for the tumor cells within the PPx cohort, informing aggressiveness and outcomes. Whether women with PPx TNBC can benefit from C3-directed therapeutics warrants further research. Further, analysis of PPx epithelia showed evidence of high metabolic turnover. Whether this high metabolic turnover in tumor epithelia is a cause or consequence of aggressive biology linked to poor prognosis warrants further analysis. Taken together, our analysis provides evidence of aggressive biology in PPx (but not GPx) non-cancerous epithelial cells. Further prospective studies are warranted to test whether expression of EMT, chemokines, and drivers of metabolic turnover, can be used to risk-stratify screen-detected pre-cancerous breast lesions.

Differential analysis of GPx vs PPx for all PanCK+ segments showed no statistically significant difference. To be able to predict the prognostic state which is agnostic to the epithelial state (J, A, I), we utilized a deep learning approach, we obtained a ranked list of genes capable of predicting the prognostic state (235 GPx and 236 PPx. The distinct ranking of several genes as identified using deep learning alludes to their differential interactions in the two groups and influence on subsequent prognosis. The features we identified are involved in MYC-centric cellular stress response; the latter has been previously implicated as a crucial determinant of tumor progression and outcomes[67,68]. Our results emphasize the complex and distinct transcriptional programs associated with epithelial state changes. Most interestingly, our findings of markers and mechanisms in atypia which posit poor prognosis combined with the detailed spatial dissection of the interactions between the tumor and the TME, provide a mechanistic basis for subsequent prognosis. Our findings have the potential to bolster and inform therapeutic considerations in women receiving treatment for localized TNBC.

## MATERIALS AND METHODS

### GeoMX Digital Spatial Profiler

Spatial transcriptomic profiling was performed using GeoMx-DSP^TM^ (DSP; Nanostring Technologies). Pre-treatment formalin-fixed paraffin-embedded (FFPE) were cut into 10 concurrent 5-mm sections. The first 5 mm section was stained with hematoxylin and eosin (H&E). The second slice was hybridized with the GeoMx Probe Mix for NGS readout (human whole transcriptome Atlas (GeoMx Hu WTA panel, Cat. #: 121401102, Nanostring Technologies) panel as per manufacturer’s protocol. The GeoMx Hu WTA panel includes 18000+ genes with several key genes involved in cancer cell biology and the tumor microenvironment. Slides were stained with GeoMx Solid Tumor TME Morphology Kit (Cat. #: 121300310, Nanostring Technologies) and GeoMx Nuclear Stain Morphology kit (Cat. #: 121300303, Nanostring Technologies) according to the manufacturer’s protocol. Regions of interest (ROI) were selected (diameter of 300 microns, minimum of 3 replicates representative of the tumor edge (including adjacent immune cells), tumor interior, and adjacent non-cancerous tissue. Pan cytokeratin (PanCK) was used to segment each ROI defining specific areas of illumination (AOI) and distinguishing the tumor (PanCK+) from the stromal (PanCK–) component. Stromal areas were further analyzed using the pan-immune marker CD45 and the pan-T-cell marker CD3. Segmentation was conducted on two classes of AOIs within each ROI: PanCK- and PanCK+. For each AOI, probes were collected by the machine and transferred to a PCR plate for library prep using Seq Code primers (GeoMx Seq Code Pack, Cat. #:121400205-121400206, Nanostring Technologies). Libraries from each AOI were pooled according to their dimension, purified by AMPure XP beads (A63880 Beckman Coulter) clean up, and resuspended in a volume of elution buffer proportional to the number of pooled AOIs. Libraries were assessed using an Agilent Bioanalyzer, then diluted to 1.6 pmol/L and sequenced (paired-end 2 x27) on Illumina NextSeq2000, with a coverage of 100 reads. FastQ files were uploaded to BaseSpace Illumina and converted into DCC files by GeoMxNGSPipeline software and utilized for all downstream processing.

### GeoMX Data QC and Processing

GeomxTools[18] was used for quality control (QC) and downstream analysis of the DCC files in R. All samples were pooled prior to QC. QC was performed in accordance with GeomxTools developer recommendations (min Nuclei = 40, min Area = 1000, LOQ= 2). Gene and sample filtering were chosen to be at 5% at which a reasonable number of AOIs and probes were retained. Samples that were marked as PTS, were extracted and re-processed independently to allow for time series analysis. In this case, the segment filtering threshold was set at 2%, while gene filtering thresholds were still maintained at 5%. The counts were normalized using Q3 normalization as recommended by GeoMx after gene filtering and transformed into log values for downstream analysis, when necessary(log_q). The comparison of interest dictated whether p-values or adjusted p-values were chosen for thresholding and identifying differentially expressed genes. Differential analysis was performed using a linear mixed model. Comparisons of interest once again dictated whether an LMM was performed with a random slope (across slides) or with a random slope and intercept (within slide).

### Differential analysis using Linear Mixed Models

The differential analysis presented here was performed using the linear mixed models (LMM) as prescribed by the GeomxTools vignette[18]. LMM allows users to account for the subsampling per tissue; in other words, adjusting for the fact that the multiple regions of interest captured in a slide/ tissue section are not independent observations. As indicated in the vignette of the GeomxTools two LMM models are possible when performing differential analysis, (i) Within-slide comparison, with a random slope: A within-slide comparison calls for comparisons of regions of interest within a slide, with each slide present in the sample set serving as a replicate. For example, in instances where we wish to assess if there exists a difference in invasive epithelia identified at the center of the tumor to the edge of the tumors, we use a within-slide analysis. In this example, we would consider only slides that have captured invasive epithelia from both center and edge. Similarly, we utilized this approach to also compare Invasive vs normal-adjacent epithelia and invasive vs atypia in both prognostic groups(ii) Across slide comparison, without a random slope. For instance, when we wish to compare all invasive epithelia coming from two distinct population cohorts say GPx and PPx. Differential analysis was performed using the entire feature space (10150) or the NMF reduced feature space as indicated in the manuscript.

### Image intensity and spatial entropy analysis

Images of each ROI and each PanCK segmented AOI region were obtained from the GeoMx DSP instrument using the “ROI Report” extraction. All images were first segmented to obtain only the portion inside the ROI region. The four-color channels were extracted, and an intensity filter was applied to remove the pixels in the lowest 10th percentile for each color channel. For the immune cell intensity analysis, the color channels within each ROI region were analyzed for the PanCK- AOI only. For each color channel, the intensity signal was computed by dividing the fraction of pass-filter pixels for the specified color by the total pass-filter pixels of all colors in the PanCK- AOI. Spatial entropy analysis was performed using the full ROI image. Using the intensity-filtered color pixels, both univariate and bivariate entropy measures were computed. For univariate entropy, we performed Batty’s spatial entropy to assess the region heterogeneity of each color channel in the ROI[19]. For bivariate entropy, we used different pairs of color channels and constructed a marked point pattern dataset for each color pair to assess the spatial relationship between the immune cells (yellow and red channels) vs epithelial cells (green channel). Spatial entropy was assessed using Altieri’s spatial entropy which was implemented in R using the SpatEntropy package[20]. We first used Leibovici’s implementation of Shannon’s relative spatial entropy to filter out ROIs where color distributions were not explained by their spatial co-occurrences (ROIs with *H*_*relative*_(*Z*) ≥ .5 were removed). For the ROIs that passed the spatial entropy filter, we then used Altieri’s spatial entropy with distances of 5,10,50,100,250 and 500 pixels which were chosen so that the distance steps would span from cell-cell interactions to half the diameter of the ROI. The spatial partial information calculated on each distance step showed a maximum value between 5-50 pixels which corresponds to the distance of ∼1 cell diameter or a cell’s nearest neighbors. The primary measure used to quantify the spatial dependence of the color pairs was the spatial mutual information, *MI*(*Z*, *W*).

Spatial entropy was assessed using the immunofluorescent image of each ROI by calculating the entropy for two bivariate distributions: CD3 vs PanCK staining and CD45 vs PanCK staining. For each bivariate distribution, both global and scale-dependent measures of entropy were calculated. The global measures, which assessed the entire ROI, included Shannon’s relative Z entropy and spatial mutual information; however, these measures are often insufficient to detect dependencies that only occur at specific length scales. Scale measures, which are assessed at different length scales within the spatial distribution, were calculated at pixel lengths of 5, 10, 50, 100, 250, and 500 and were chosen to span distances from cell neighbors (∼15μm) to the radius of the ROIs (∼150μm). We used Altieri’s spatial entropy to produce both spatial mutual information and scale metrics from which we used the spatial partial information to assess the dependence of the two channels[21]. For each immunofluorescent ROI image, the relevant channels were extracted, and an intensity filter was applied as previously described. The pixel locations from the two channels being assessed were converted to a point pattern dataset and the entropy measures were calculated using the relevant functions from the R SpatEntropy package[20]. To compare the spatial entropy metrics between different sample classifications, a two-sided T-test was performed, and the FDR p-value was assessed for significance at the 0.05 level.

### Non-negative matrix factorization (NMF) analysis

Non-negative matrix factorization (NMF) is a powerful dimensionality reduction and clustering technique and has proven particularly useful for identifying underlying patterns and structures in high-dimensional non-negative gene expression data. In this study, we applied NMF using the NMF library in R[22], on log_q data generated from PanCK+ segments in GPx and PPx (to mitigate the influence of extreme values and provide a more stable representation of the underlying biology).To ensure the robustness of our NMF model, we assessed its stability using a consensus matrix approach. Specifically, we set the number of runs (nrun) to 5 and tested ranks ranging from 2 to 10. This process was iterated 20 times with different random seeds. The best rank for each seed was determined based on metrics such as cophenetic correlation, dispersion, and silhouette score, ensuring a comprehensive evaluation of model performance. Subsequently, the selected best rank for each random seed was utilized to run the NMF model for 100 iterations (nrun = 100), allowing for further refinement and convergence of the factorization process. We extracted the metagenes using the "extractFeature" function with the method parameter set to "combine" within the NMF library. These metagenes represent a compact and interpretable representation of the original features, capturing the consensus patterns across multiple runs and random seeds. Finally, the union of all 20 seed runs was utilized as the reduced feature space, referred to as “consensus features” for downstream analysis. These consensus features encapsulate the most relevant and stable biological signals present in the dataset, enabling effective interpretation and subsequent analyses.

### Spatial cell-type deconvolution using SpatialDecon, and a TNBC-specific reference matrix

The reference matrix readily available for deconvolution of the TME within SpatialDecon[22] v1.1(called SafeTME) is largely derived from the immune and stromal cells derived from PBMCs and flow-sorted tumor cells, we chose to develop a TNBC-specific deconvolution matrix using the “create_profile_matrix” function of the SpatialDecon package. This function requires a cell * gene count matrix, along with the cell annotations as data input. To achieve this, we downloaded the scRNAseq data from a recently published study specific to breast cancers[11] (downloaded from https://singlecell.broadinstitute.org/single_cell/study/SCP1039/a-single-cell-and-spatially-resolved-atlas-of-human-breast-cancers#study-download) containing 100064 cells. From this, we extracted all single cells from all TNBC (42512 cells). Since we were specifically interested in generating a reference matrix for deconvoluting the PanCK- segments of the ROIs, we excluded epithelial cells (cancer and normal) bringing down the # of single cells used from this study to 30719. As provided in the supplementary of the original study, we retained the cellType_major and cellType_minor information across 30719 cells. We focused on 7 major types as captured by the cellType_major information, including B-cells, T-cells, Endothelial, Plasmablasts, perivascular-like (PVL) (with increased expression of MCAM/CD146, ACTA2 and PDGFRB), Myeloid, and CAFs (renamed as Fibroblast/Mesenchymal)) to drive the generation of our reference matrix. As previously described[23], we further processed and filtered these ∼30K cells using Seurat[24] v4.3.0.1, resulting in a filtered data matrix with 11951 features x 26451 cells. The TNBC specific reference profile matrix using the “create_profile_matrix” command (minCellNum=10, scalingFactor=1 and minGenes=100) in SpatialDecon v1.1, with the expression of 1066/11951 genes identified as breast cancer-specific serving as input. This reduced gene list was obtained by amalgamating the genes in the safeTME matrix, the genes expressed in breast TME as published in Navin et al[25], and 135 breast-specific genes as published in the Human Protein Atlas (https://www.proteinatlas.org/humanproteome/tissue/breast), belonging to the 7 major cell types. SpatialDecon v1.1 was subsequently also utilized to generate the spatial deconvolution of the TME using the “RunSpatialDecon” function and the reference matrix generated above as profile matrix. Proportions of cell counts as well as the abundance estimates were extracted and plotted, as outlined in the vignette.

### EcoTyper

EcoTyper was implemented using their source code available at [https://github.com/digitalcytometry/EcoTyper accessed Jan 2nd, 2024]. TPM transformed filtered probe data (before Q3 normalization) for each segment and served as input to EcoTyper. The cell states were computed using the ecotyper_recovery_bulk.R script, following their vignettes. Given the nature of the GeoMX data, we further multiplied the cell state abundances determined by EcoTyper, with the cell type abundances for each of the overlapping cell types to get a final cell state abundance for each AOI. AOIs were assigned to the cell state if the state exceeded the 80^th^ quantile value of the final cell state abundance computed for each cell type. The description of the cell states was obtained from supplementary table 4 of the original publication[26], and the set of all cell state markers used for generating the violin plots, was obtained from the authors of the original publication (personal communication).

### BulkSignalR

BulkSignalR[27] was utilized to infer the ligand-receptor interactions (LRI) in the TME of the Tenr subset. BulkSignalR utilizes a curated database of 3249 ligands, receptors, their associated pathways and downstream target genes as the background from which to make the inferences. For the L-R interactions captured in the TME, Q3-normalized data from all 10150 features across 61 GPx Tenr AOIs and 40 PPx Tenr was utilized in this analysis (set. Seed(123), min.count = 0 and prop = 0). For the L-R interactions described in atypia, we used Q3-normalized data from all 10150 features, in the ROIs with atypic epithelia. We included LRI pathways with a minimum of 4 downstream target genes present in our gene list (min.positive = 4). BulkSignalR requires null distributions of Spearman correlation coefficients between a ligand and receptor (L-R) and receptor and its target genes (R-T). The Gaussian kernel-based empirical model was identified as the appropriate statistical model for L-R and R-T correlations in both GPx and PPx. When inferring significant LRI, those with L-R correlation > 0.25 and association with a pathway with q < 0.05 were included in the final list for GPx and PPx. Pathway significance was calculated as a combination of L-R and R-T correlation significance, and the most significant pathway for each unique L-R was used.

### Enrichment and TF target analysis

ssGSEA was applied to expression data from each AOI, separately for the PanCK+ and PanCK- segments, using the Hallmark gene set from MSigDB using the GenePattern software[28]. Additionally, all enrichment presented in this manuscript were performed and plotted using either clusterProfiler[29] or enrichR[30]. TF target analysis was performed using DecoupleR[31]. String-db v12 was utilized to identify protein-protein interactions (medium confidence) in gene sets of interest. All the networks were then exported to Cystoscape and visualized in Cystoscape[32]. Enrichment of the String PPI was performed within Cystoscape using GO_BP, Wiki pathways, and Reactome pathways categories. The enrichment reported was identified with a fdr<10^-4^ or better.

### Convolution neural networks

NMF-reduced gene expression data from pooled GPx and PPx AOIs was utilized to identify a gene signature defining the epithelial state for each prognostic group, using convolution neural networks [previously published]. We split the data into distinct subsets, consisting of training, validation, and testing sets using a stratified splitting strategy. The use of such a strategy guaranteed a balanced representation of GPx and PPx classes in each group. Specifically, we allocated 10% of the data for testing, 18% for validation, and reserved the remaining 72% for training. Furthermore, to ensure reproducibility, we incorporated a random seed mechanism during the allocation of data, ranging from 1 to 500, facilitating consistent data partitioning across iterations. Once the data was appropriately partitioned, we reshaped each sample’s original 1D structure to 2D. We instantiated our CNN with this restructured data to allow for the use of traditional 2D CNN layers. The Borda count method was utilized for ranking genes/features as obtained from our saliency maps. Borda count is a ranked polling method that takes the position of each element and assigns a value for its corresponding position. Elements lower in ranking receive fewer “points” and are ranked less highly in the aggregate ranking. We assessed the concordance between the two ranked lists generated, utilizing the *aggregateRanks* function (method= “Stuart”) available via the RobustRankAggreg[33] v1.2.1 package in R. Genes with a score of 0.05 were identified as similarly ranked, while genes with a score >0.5 were identified as being most dissimilar between the two gene lists

## List of Supplementary Materials

Data file S1- Containing supplementary tables S1-S10

Fig S1-S6 Supplementary figures

## Acknowledgements

The authors wish to thank Professor Jason Swedlow and Dr. Josh Titlow from the Wellcome Leap Foundation for invaluable discussions.

## Author contributions

K.M and S.S designed the systems biology analyses, with K.M. performing majority of analyses presented within the paper. D.V analyzed the ligand-receptor interactions and provided support to K.M. with the analyses. D.F performed all the necessary processing and analysis of the image files. S.A designed and implemented the CNN to classify patient groups. Z.M generated the molecular subtyping of TNBC utilized in this study. L.Y, J.T and R.P designed the GeoMX assays and generated the data. D.S and V.S were lead pathologists in this study, with V.S identifying and annotating all ROIs considered within this study. X.C.W, M-A L.B identified and selected the tumors and their respective metadata from the LSU tumor registry. L.M and A.O were involved in fine tuning the patient cohort based on prognosis, used in this study. S.S. designed and supervised the entire study with support from V.S. K.M. wrote the first draft of the manuscript, with revisions by S.S. All authors have read and agreed to the submitted version of the manuscript.

## Data and materials availability

The 1440 DCC files generated and utilized in the study are available to reviewers at www.tnbcworkbench.org/tnbc_respository.php. Due to the nature of the data, this is currently controlled access and will be made open source after peer review. All other additional material necessary to recapitulate the results are provided in the supplementary materials attached. The code generated in this study largely follows the open-source vignettes associated with each R/BioC library but can be made available upon request.

---

**Fig. S1.**
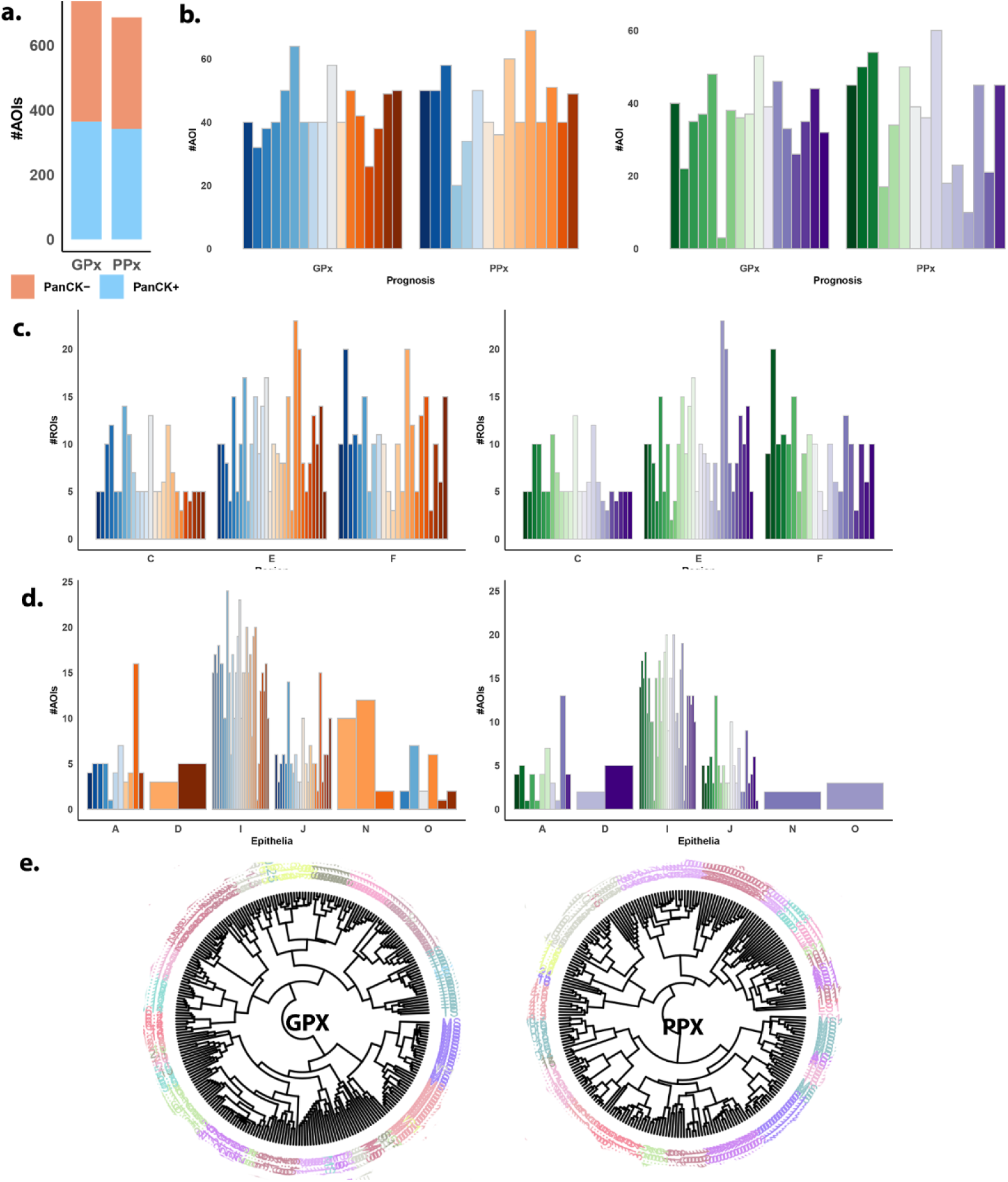
a. A table showing the S-E-R- L-P-D-N annotation schema developed in this study. b. Barplot showing the distribution of total AOIs captured pre-processing (1424 AOIs - 73 7 GPx and 687 PPx) c. the distribution of AOIs pre-processing (left panel) and post-processing for both GPx and PPx samples. Each bar represents a patient sample d. Frequency of the AOIs captured in the various regions (Center-C, Edge-E and Isolated foci- F) of the tumor FFPE slice, pre (left panel) and post-processing (right) across both GPx and PPx samples. Each bar represents a patient sample. e. Frequency of the AOIs captured with various epithelial types as annotated by the pathologist in accordance with extended figure 1a., pre (left panel) and post-processing (right) across both GPx and PPx samples. Each bar represents a patient sample. f. A circle dendrogram capturing the heterogeneity (intra and inter-patient) of the expression within ROIs. Every patient is represented by a unique color

**Fig. S2.**
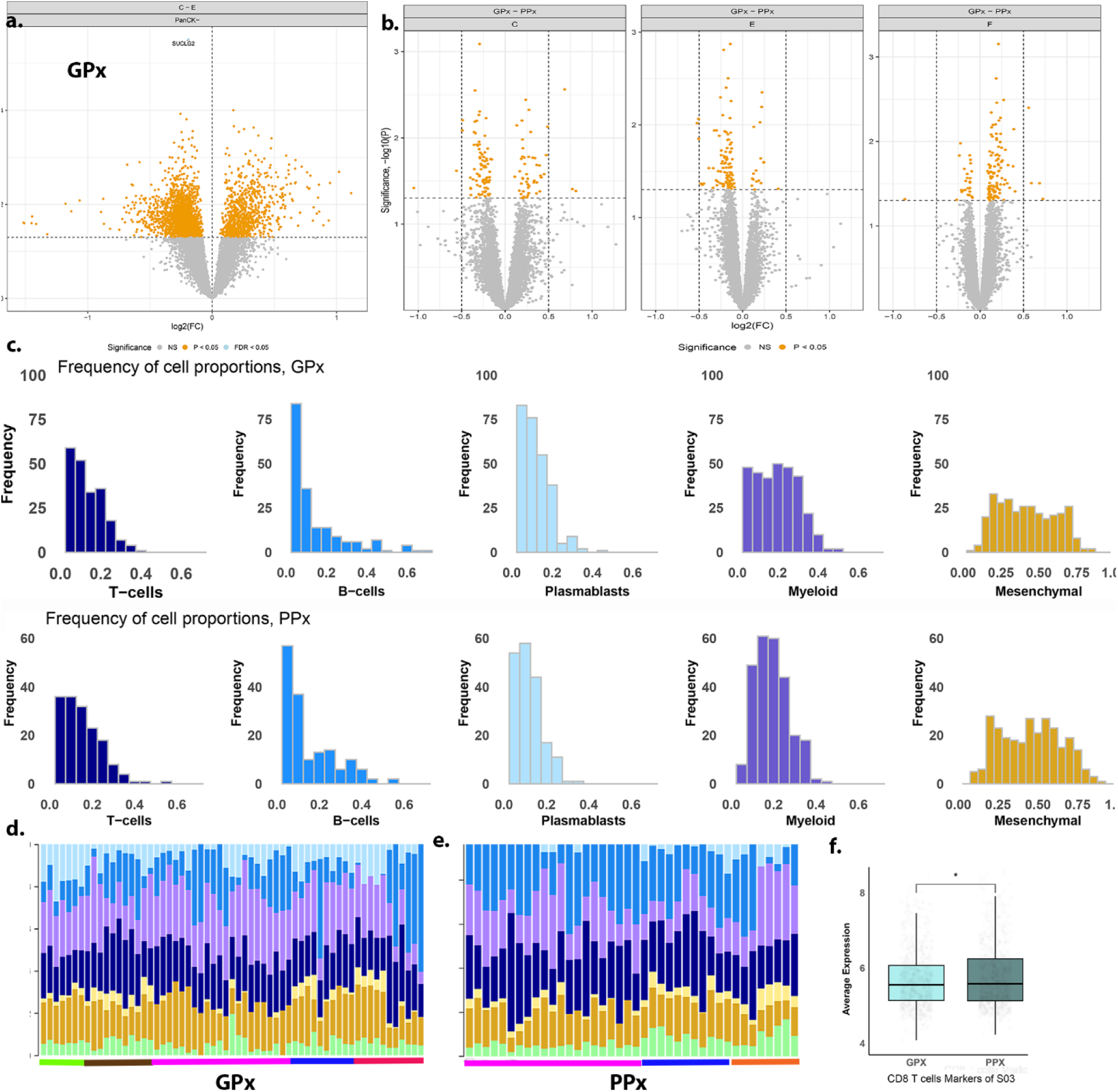
a. Differential expression captured in the PanCK- segment of GPx, between center and edge AOIs. b. Differential expression for GPx vs PPx between AOIs captured in each region (C-center, E-edge and F-isolated foci) of the tissue. c(top and bottom) captures the cell type frequencies of the T-cells, B- cells, Plasmablasts, Myeloid cells and mesenchymal (PVL+fibroblasts) in GPx and PPx PanCK- AOIs respectively. d. The abundances of the seven cell types estimated by spatial deconvolution the 61 AOIs from 5 GPx patients. Each color bar represents a patient. e. The abundances of the seven cell-types estimated by spatial deconvolution the 40 AOIs from 3 PPx patients. Each color bar represents a patientf. Average expression of CD8T cell markers in S03 (exhausted T) shows a significant (p<0.05) increase in PPx - Tenr AOIs over GPx- Tenr AOIs

**Fig. S3.**
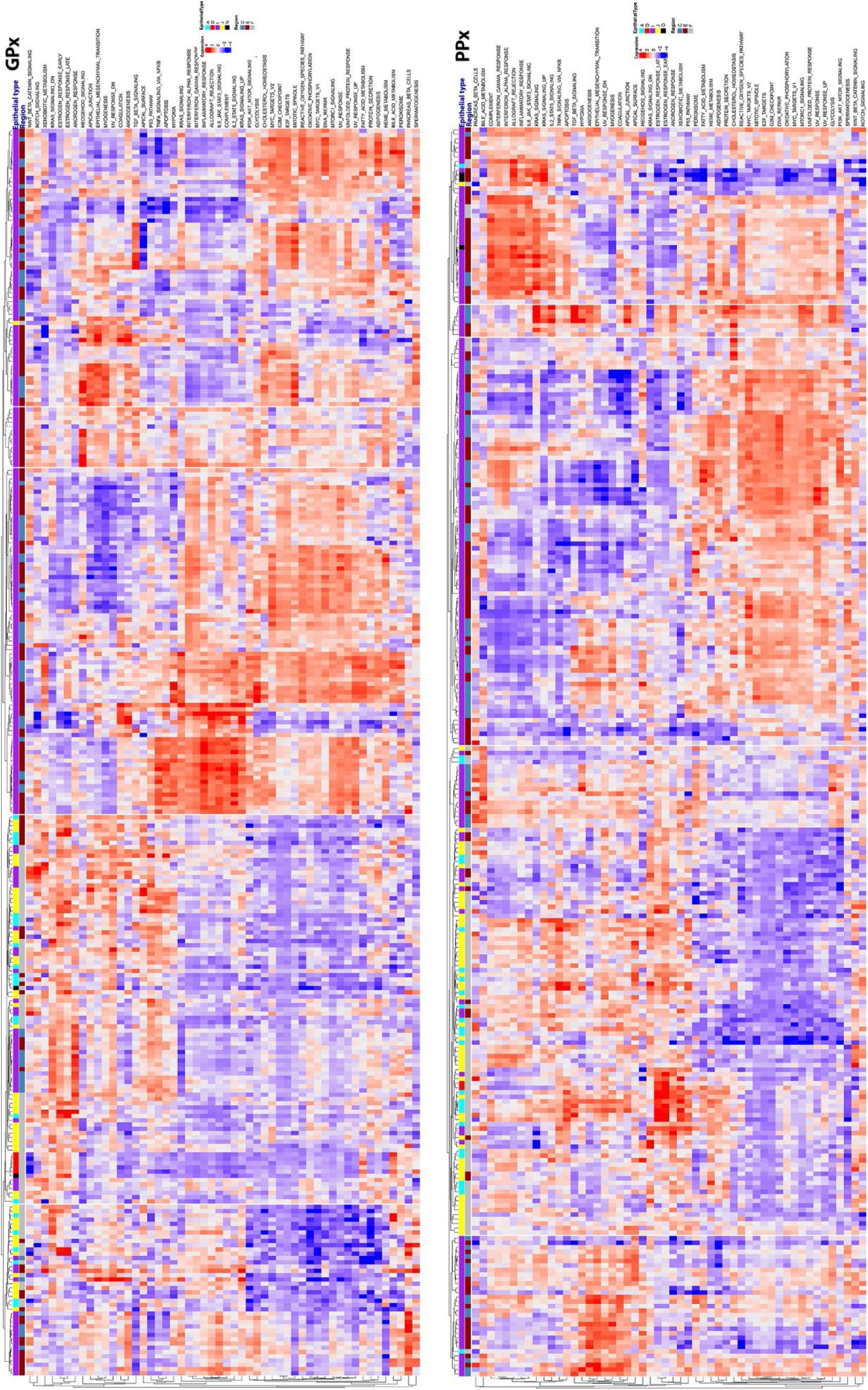
ssGSEA on PPx and GPx epithelia (PanCK+ AOIs) show an ssGSEA results highlighted a prominent increase in mechanisms associated with proliferation, DNA repair and metabolism within the invasive epithelia, in both GPx and PPx The AOIs with non-invasive epithelia in both subsets were more prominently associated with signaling cascades in volved in in EMT including WNT Beta catenin, Notch, and KRAS signaling.

**Figure S4.**
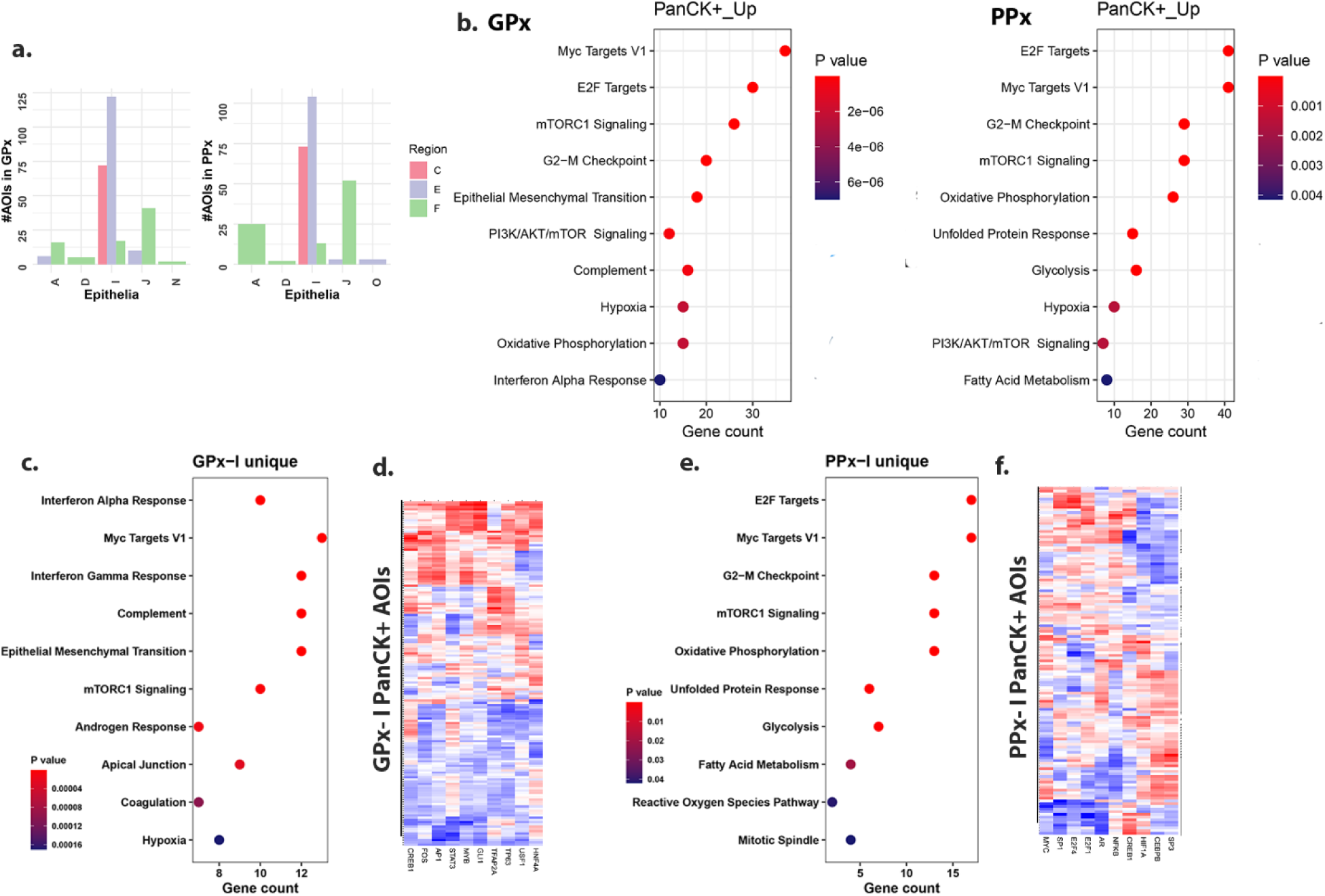
a. Frequencies of AOIs with each epithelial type captured in the diAfferent regions within GPx andPPx. b. Enrichment (Hallmark gene sets) of the genes upregulated within AOIs within invasive epithelia compared to the normal adjacent in GPx(left) andPPx (rightpanel). c. Enrichment of genes uniquely upregulated in PanCK+ AOIs (left) and TF target enrichment as identified by DecoupleR includingSTAT3, API TFs, and TP63. d. A similar analysis on the DEGs unique in PPx AOIs with invasive epithelia, with enrichment ofMYC, E2F, H1F1A and NFKB transcription factors

**Figure S5.**
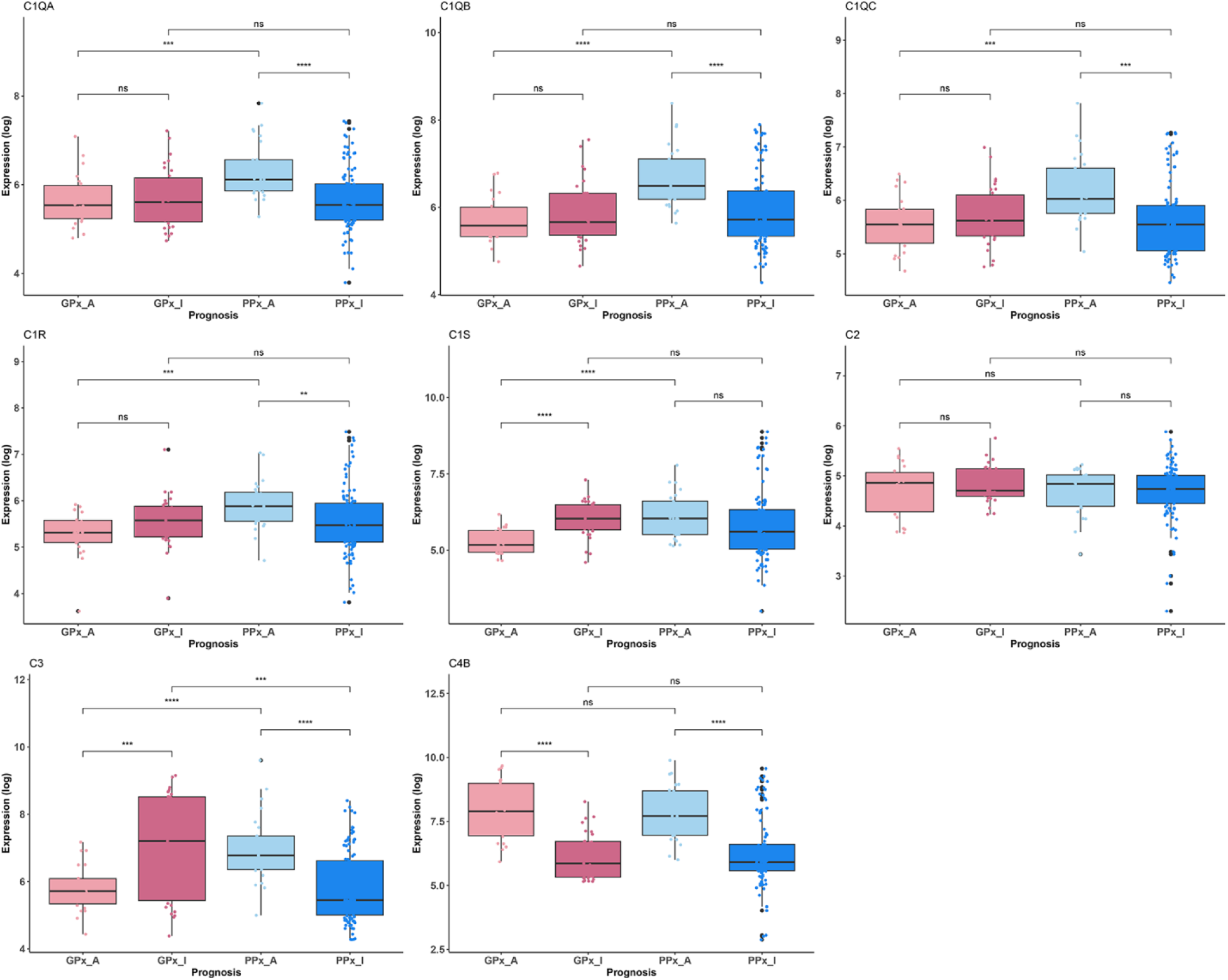
Boxplots showing expression(log2) of complement genes in GPx and PPx PanCK+ AOIs with atypia and invasive types.

**Figure S6.**
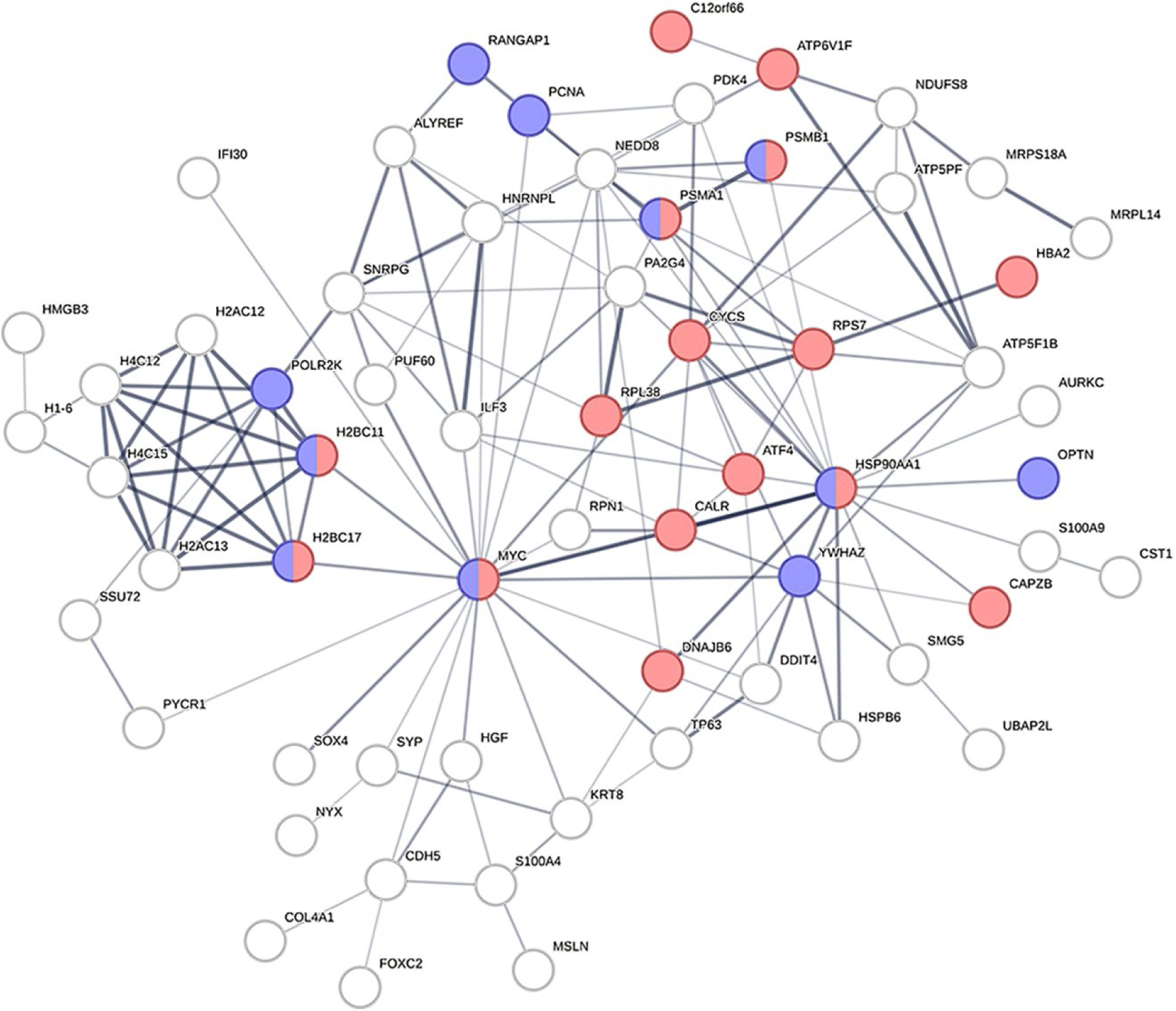
This network represents the protein-protein interactions captured between genes that were identifed to be highly dissimilar between the two ranked feature lists by our CNN as necessary to predict the labels GPx and PPx. The enrichment of this networks highlights that the genes are mostly associated with cellular stress response (red) and cell cycle control/DNA damage repair (purple) centered on MYC emphasizing the difference in the interactions that exist in epithelia which may contribute to subsequent prognostication.

